# Bats from the Colombian Caribbean Reveal a new subtype of Influenza A (H18N12)

**DOI:** 10.1101/2024.08.25.609627

**Authors:** Daniel Echeverri-De la Hoz, Caty Martínez-Bravo, Bertha Gastelbondo-Pastrana, Ricardo Rivero, Yesica Lopez, Richard Hoyos, Maira Alemán, Evelin Garay, Valeria Bertel, Germán Arrieta, Juan David Ramírez, Salim Mattar

## Abstract

Influenza viruses have an excellent capacity for mutation and adaptation in mammalian hosts, which makes them viruses of medical and veterinary importance. Influenzaviruses have been studied mainly in birds but minor in bats. It is unknown whether Chiroptera are reservoirs of influenza viruses. However, circulation in bats showed molecular divergence from H17N10 (Guatemala) and H18N11 (Peru), and they were designated as new subtypes. The study aimed to characterize the influenza A virus detected in the fishing bat *Noctilio albiventris*. A surveillance study of pathogens of public health interest was carried out; rectal samples were taken from four fishing bats (*N. albiventris*) captured in Talaigua Nuevo, Bolívar, Colombia. The samples were sequenced by NGS using DNBseq (MGI-G50®) and analyzed with bioinformatics tools. Eight viral contigs associated with the Orthomyxoviridae family were obtained. The identified segments showed around 90% similarity with H18N11, except for the neuraminidase (N). The phylogenetic analysis of the N protein showed the appearance of a basal branch to the N11 subtype, and the molecular clock indicates that it does not share a recent common ancestor. 3D modeling indicates that the N protein of *N. albiventris* presents three mutations (K363R, T242K, and I139V) near the hypothetical active site of the protein. These mutations potentially increase the interaction with the HLA-DR of bats, which could have significant implications for the virus’s behavior. The phylogenetic, evolutionary, and antigenic divergence of the N protein of *N. albiventris* suggests a new subtype called H18N12. Its role as a pathogen must be studied.

## 1 Introduction

Bats act as sentinels of epidemiological surveillance because they are hosts of viruses that cause important diseases such as coronaviruses, paramyxoviruses and filoviruses [1]. In addition, Venezuelan Equine Encephalitis [2], and dengue virus [3, 4]. Although bats have not yet been established as reservoirs of influenza viruses, a new genomic sequence of influenza A virus (IAV) designated H17N10 was detected in fruit bats in Guatemala [5]. Later, another genome classified as H18N11 was characterized in bats in Peru [6]. Also, a virus similar to avian strains was detected in Egyptian bats [7]. These findings allow us to think about new hypotheses of the origin of IAVs and their impact on public health [5, 6].

The origin of bat IAVs needs to be better known, and few studies have clarified their origin. Phylogenetic and evolutionary analyses established that they come from a common ancestor with the avian IAV subtypes [8]. However, divergence is one of the features that has drawn the attention of researchers. IAVs probably split into two genetic branches due to geographic separation and multiple early spread events, or IAVs may have undergone drastic changes to adapt to bats [9].

Crystallographic structures of the hemagglutinin (H) and neuraminidase (N) proteins have been generated to characterize bat IAVs [6, 10, 11]. The structures were similar to avian IAV strains but with specific molecular modifications as an adaptability mechanism to bats. In that sense, IAVs do not use sialic acid but molecules of the major histocompatibility complex class II (MHC-II) as cell entry receptors [12, 13]. This mechanism suggests adaptability and possible inter-species jumping since MHC-II is expressed in immune system cells and epithelial tissue in many animals, including pigs, mice, and chickens [9].

On the other hand, the N protein of bat IAV appears to have no catalytic activity, and its function is an enigma for researchers. The results indicate that the N protein could induce a low expression of MHC-II molecules by an unknown mechanism [9]. If these data are confirmed, it would demonstrate that the surface glycoproteins of bat IAVs have receptor binding and destruction activities. Therefore, bat IAVs would carry out the same infection and release process of viral particles as avian IAVs [8]. However, these hypotheses have yet to be corroborated.

The objective of the present work was to characterize a new sequence of Influenza A viruses detected in the fishing bat *Noctilio albiventris* from Colombia.

## 2 Results

### Genomic, phylogenetic, and evolutionary analysis of influenza A virus detected in *N. albiventris*

Initially, three contigs assigned to the H18N11 subtype were obtained. The reference genome map yielded seven segments corresponding to the PB1, PB2, PA, HA, NS, NP, and M genes. The search effort for the NA gene was increased by decreasing the alignment and similarity criteria at the seed level. Eight segments were obtained that corresponded to 0.12% (22,256/18’129,755) of the reads with a depth ¿ 100X. The genome comparison analysis shows greater similarity with the H18N11 subtype sequence recorded in Peru (93%). The percentage of similarity of each segment varied between 92 and 98%, and the N protein gene of *N. albiventris* showed greater divergence (Table 1). Figure 2 shows the identity patterns between the sequence detected in *N. albiventris* concerning the H1N1, H2N2, H3N3, H17N10, and H18N11 subtypes. Phylogenetic analysis indicates that the HA segment is closely related to node H18 from Peru, Brazil, and Bolivia (Figure 3A). However, the NA segment forms a basal branch (Figure 3B). Based on phylogenetic analysis and N protein divergence, evolutionary time tracing indicates that TMCRA gave rise to the bat NA segment in 1420, and the *N. albiventris* NA segment comes from a node in 1941 (Figure 3C). This demonstrates that the previously described N11 segment is phylogenetically related to *N. albiventris* NA but does not share a recent common ancestor.

**Table 1.** Percentages of similarity of influenza A virus/*N. albiventris* (Colombia) segments concerning the H18N11 subtypes obtained in Peru and Bolivia.

| Gene | H18N11-Peru |  | H18N11-Bolivia |  |
| --- | --- | --- | --- | --- |
|  | Amino acids | Nucleotides | Amino acids | Nucleotides |
| PB2 | 98.68% | 92.67% | 98.12% | 92.32% |
| PB1 | 99.6% | 95.28% | 98.81% | 94.97% |
| PA | 99.86% | 97.31% | 99.58% | 97.27% |
| HA | 100% | 98.74% | 100% | 96.21% |
| NP | 99.80% | 94.73% | 100% | 98.49% |
| NA | 92.39% | 81% | 92.17% | 80% |
| M | 100% | 98.81% | 100% | 91.03% |
| NS | 99.55% | 96.35% | 100% | 98.45% |

### Antigenic structure and molecular docking of the N protein

Amino acid sequence analysis shows a domain conserved at the level of the *Alphainfluenzavirus* genus and another of sialidase enzymes. The 3D structure for the N protein was generated from the crystallographic structure of the N11 protein of the H18N11 subtype from Peru (4mc7.1.A). Computational modeling generated a tetramer structurally similar to the N protein of avian strains that contains six antiparallel *β*-sheets in a helix-like arrangement. It also maintains six residues (R118, W178, S179, R224, E276 and E425) conserved in the active site. The alignment with the H18N11 subtype from Peru and Bolivia, has yielded points of divergence at the level of the transmembrane and distal domains of the protein (Figure 4A). The most significant mutations are highlighted in the hypothetical active site region (Figure 4B). At the conformational level, our analysis of the crystal structure of the N11 protein from Peru (4mc7.1.A) and the *N. albiventris* N protein has revealed critical structural differences. The hypothetical active site pocket of the N11 protein is broader than that of the IAV and influenza B virus N proteins due to the movements of loops 150 and 430 [6] (Figure 5A). Conversely, the hypothetical active site pocket of the *N. albiventris* N protein is narrow due to the plasticity of loop 150 and the K363R mutation (Figure 5B). This unique structural feature has led to a statistically significant interaction between the *N. albiventris* N protein and HLA-DR (MCH-II) of bats (8JRJ) (Figure 5C), with nine residues involved in hydrogen bond formation being identified (Figure 5D). Notably, three of the five mutations (Ser361, Ar363, and Lys242) have been found to increase the binding with the S1 subunit (110-165) of the *α*2 chain of bat HLA-DR, forming bonds with high specificity and strength (Å2¿1500). The movement of loop 150, which reduces the space of the active site, allows for a more significant contact and interaction surface. These findings have significant implications for our understanding of viral protein interactions and could potentially inform future research and drug development.

## 3 Methods

### Type of study, location and ethical aspects

A prospective, cross-sectional descriptive study was conducted with rectal swab samples collected from bats captured between January and December 2023 in Talaigua Nuevo, Bolívar, Colombia (Figure 1). The collected samples are part of an epidemiological surveillance study of emerging viruses in bats and mosquitoes developed by the University of Córdoba, Colombia. The Institute of Biological Research of the Tropics and ethics committee of the University of Córdoba approved the project under the permission of the National Environmental Licensing Authority of Colombia (Resolution 00914 of August 4, 2017). All animals captured in the study were released.

**Fig. 1.**
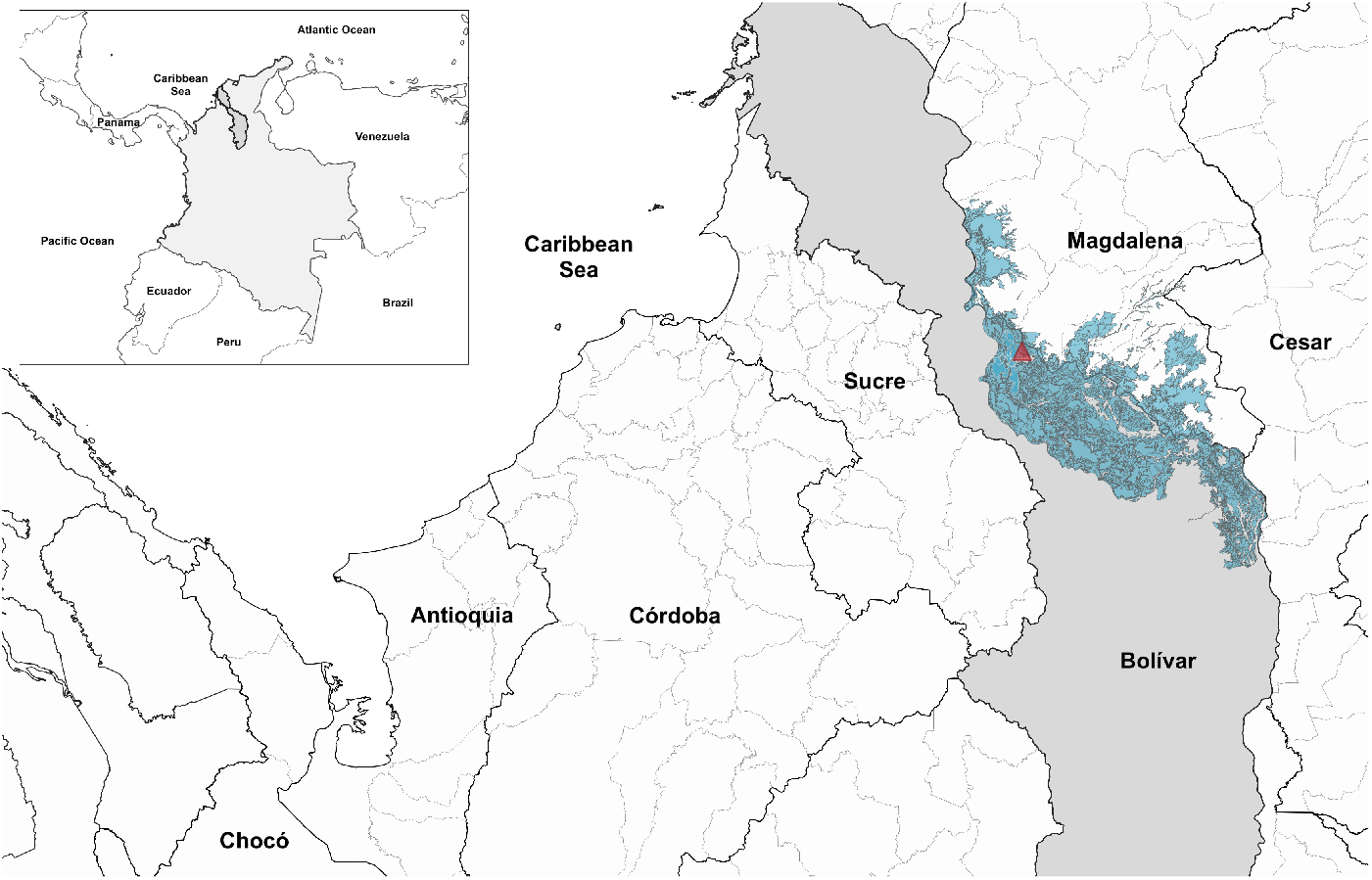
Geographic location of the department of Bolívar, Colombia. The red triangle shows the sampling point (9°18’28”N-74°36’56”O).

**Fig. 2.**
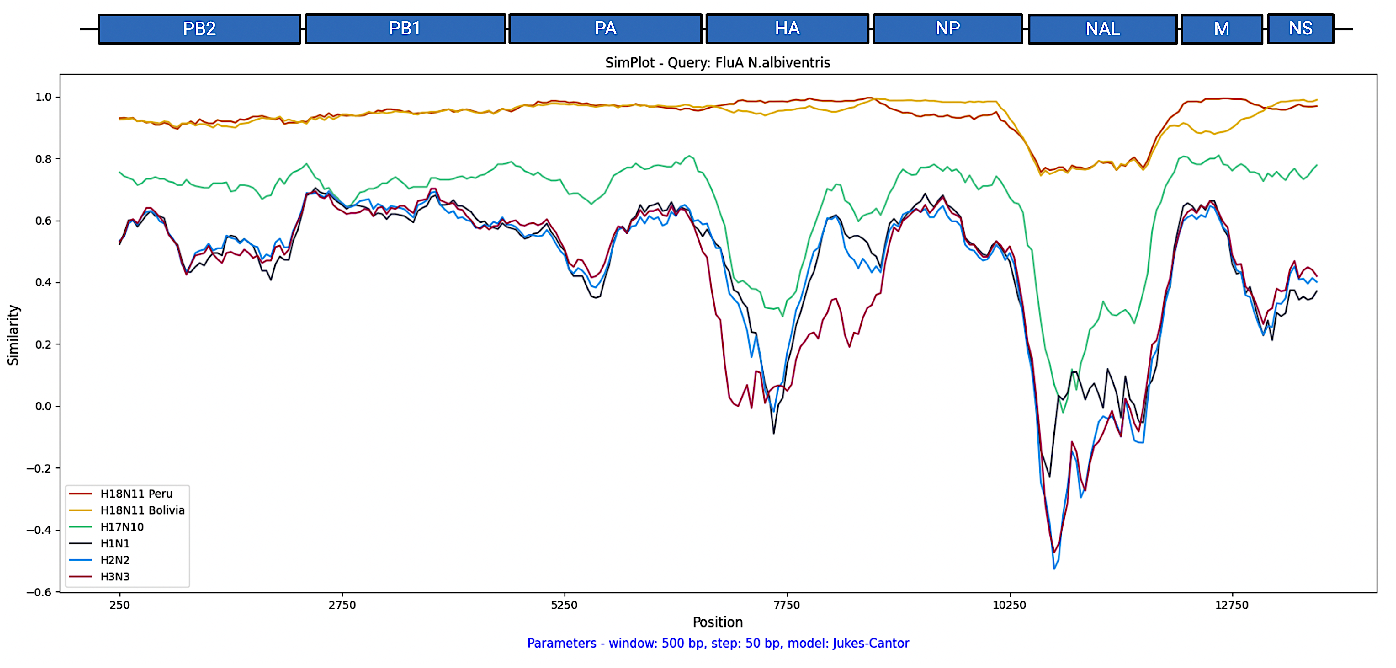
Identity patterns between the sequence detected in *N. albiventris* concerning the H1N1, H2N2, H3N3, H17N10, and H18N11 subtypes. The comparison was made with the amino acid sequences for each segment. A 500bp visualization window with 50bp steps and Kimura as a substitution model was established

**Fig. 3.**
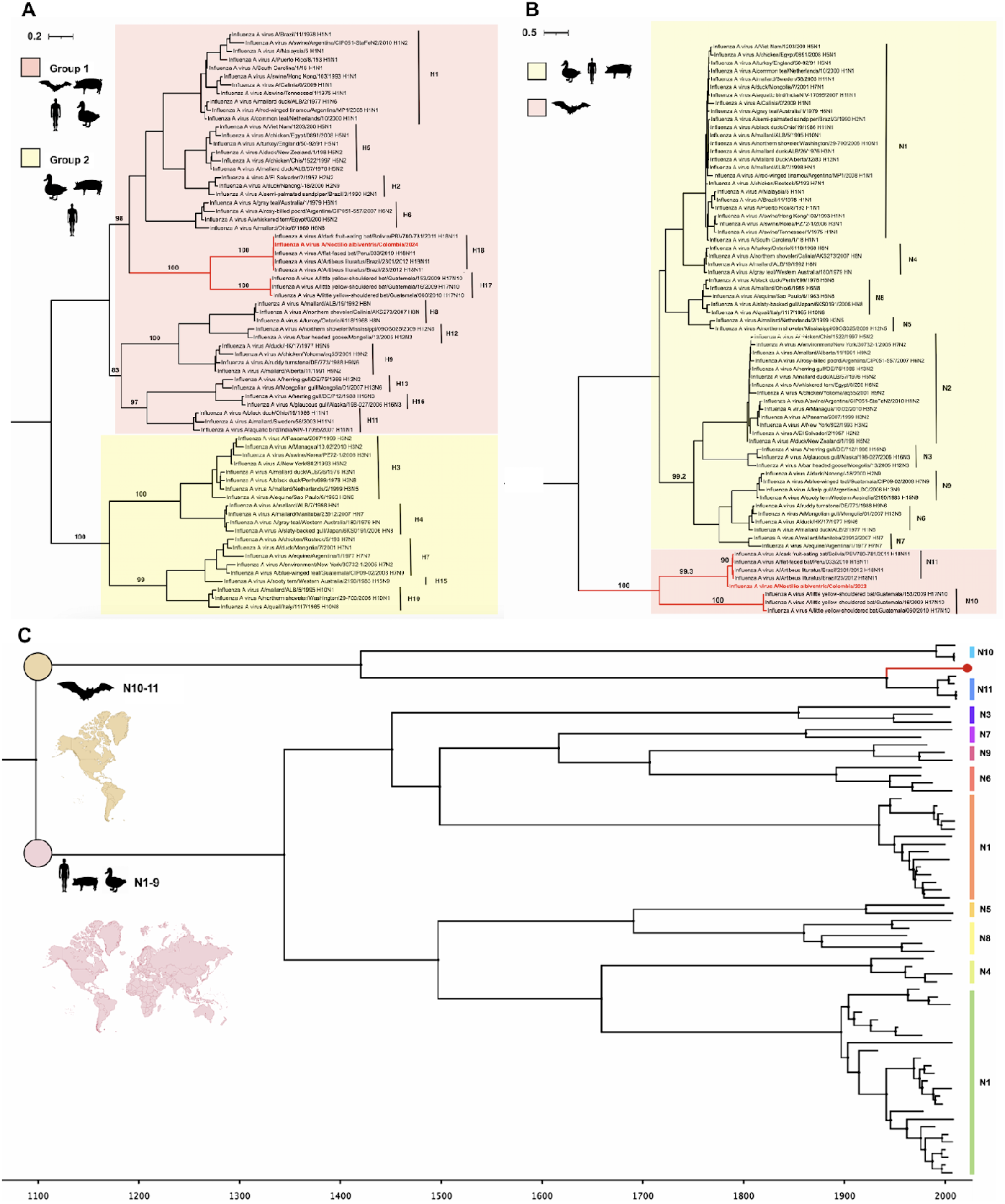
Phylogenetic tree of the influenza A virus/*N. albiventris* HA (A) and NA (B) genes. Bat influenza A branches and the new sequence are presented in red, which signifies their unique evolutionary path. The phylogenetic tree was constructed using the maximum likelihood method. Bootstrap values (1000) were given at the relevant nodes. The branch scale represents the number of amino acid substitutions along the tree branches. Reference sequences from all reported IAV subtypes (HA1-16 and NA1-9) were included in all trees. C. Molecular clock to determine the TMCRA of the NA segment. The branch of the NA segment of *N. albiventris* is represented in red and arises from a node dating back to 1940. The node that gave rise to the N11 subtype dates back to around 1990. A chain length of 100,000,000 interactions and a sampling frequency 10,000 were established. The scale represents the estimated time from the common ancestor to the current sequences.

**Fig. 4.**
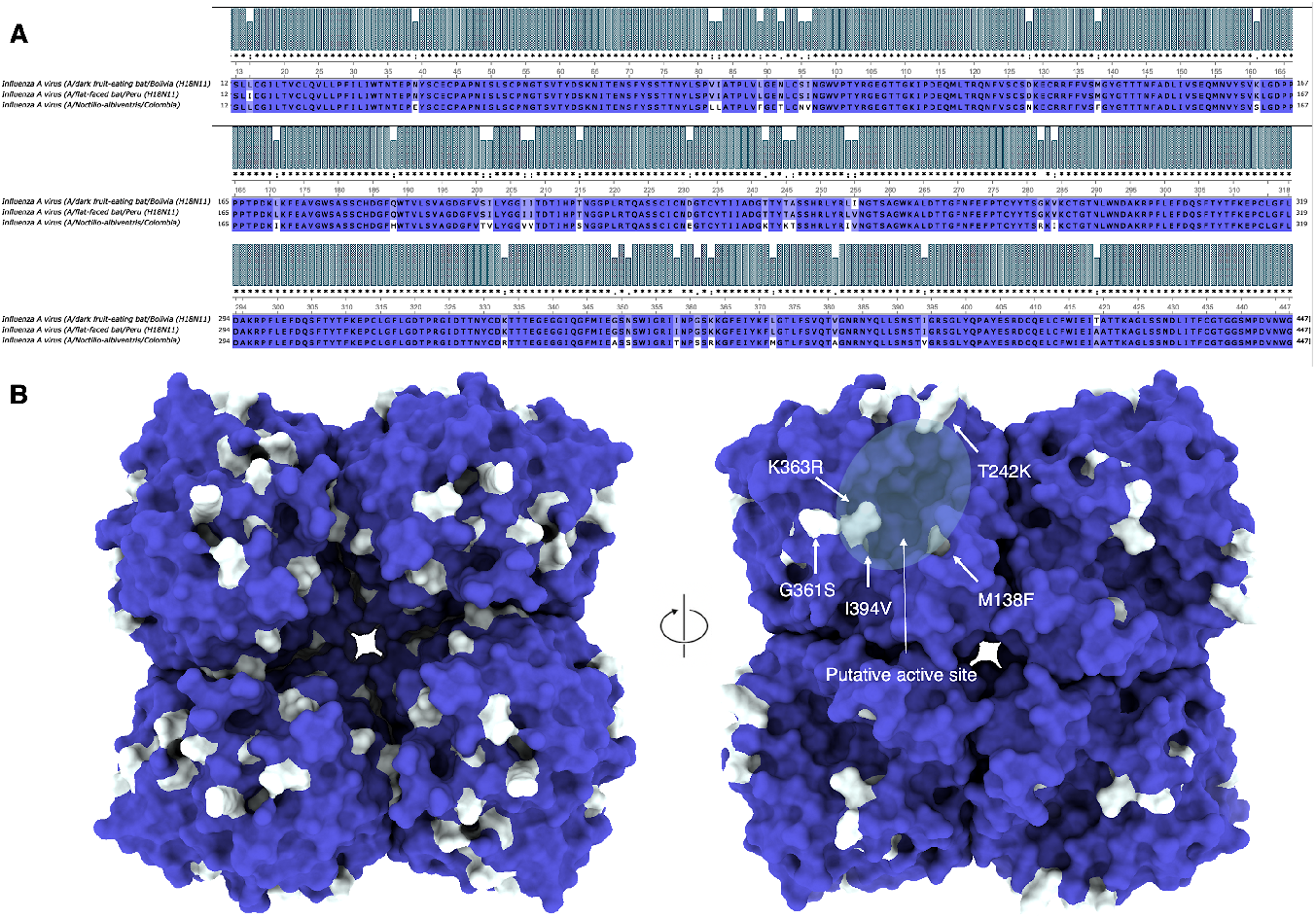
Homology modeling of the *N. albiventris* N protein. A. Alignment of the amino acid sequence of the *N. albiventris* NA protein with the N11 subtypes from Peru and Bolivia. B. 3D structure of the tetramer from a bottom transmembrane view and a top view showing the putative active site of the protein with five mutations (K363R, T242K, M138F, G361S, and I139V) relative to the N11 sequence from Peru and Bolivia. Conserved regions are shown in purple, and divergent amino acids in white.

**Fig. 5.**
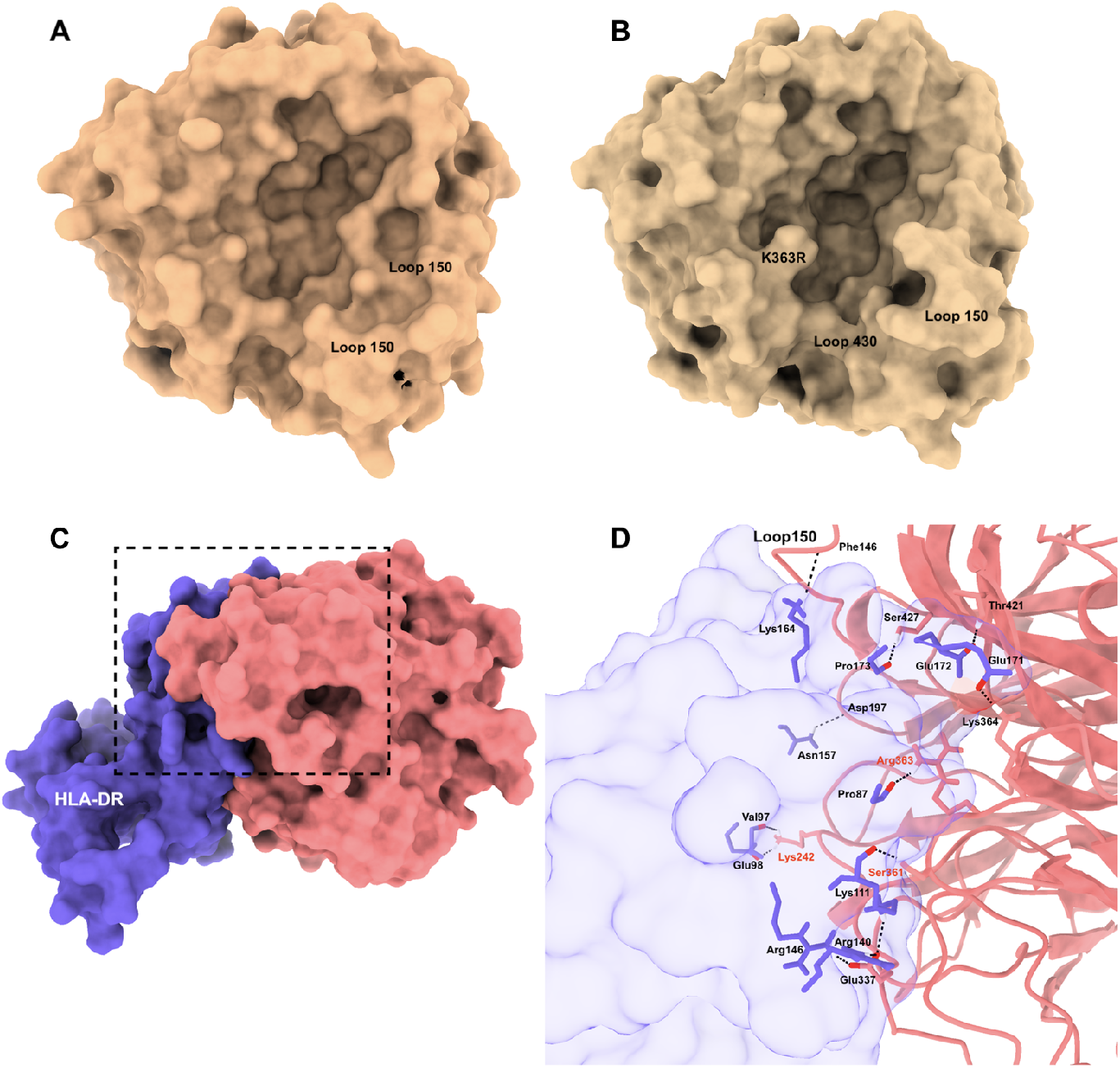
Molecular analysis and docking of the N protein. A A/bat/Peru N11 protein. B. N protein of *N. albiventris*. C. Monomer of the N protein of *N. albiventris* bound to the bat HLA-DR. The contact region (2935.1 Å2) between the two proteins is indicated. The Van der Waal energy was - 67.8. D. Binding details between residues of the N protein of *N. albiventris* and bat HLA-DR. The binding of the three mutations Ser361, Ar363, and Lys242 of the hypothetical active site of the N protein with Lys111, pro87, and Val97/Glu98, respectively, with the bat HLA-DR is evident.

### Capture of bats and sample collection

The capture was done through mist nets (6×2m) located near the water bodies of the municipality of Talaigua Nuevo. The animals were taxonomically identified using dichotomous keys based on morphometry [14]. Then, rectal swab samples were taken from four fishing bats (*N. albiventris*), which were placed in a viral transport medium stored in N2 and transported to the Institute of Biological Research of the Tropics of the University of Córdoba, where they were kept at −80°C until processing.

### RNA extraction, purification, and sequencing

Rectal samples were vortexed for 30s, and then a pool was made with each supernatant of the four *N. albiventris* samples. RNA extraction was done from 200 *µ*l of the supernatant using the GeneJET Viral DNA/RNA Purification Kit (Thermo Fisher Scientific™). The RNA was subjected to degradation of contaminating DNA with DNAse I (Promega™). Then, it was purified and concentrated with the GeneJET RNA Cleanup and Concentration Kit (Thermo Fisher Scientific™). The concentration and integrity of the RNA was determined by fluorometry with Qubit® (Thermo Fisher Scientific™). Finally, they were processed with the Paired-End FCL 150 MGIEasy Fast RNA Library Prep Set™ under the high-throughput sequencing methodology based on DNA nanobeads (DNB)from MGI Tech™. Metatranscriptomic sequencing was performed on the MGI-G50® (Shenzhen, China).

### Bioinformatics analysis

The sequences were subjected to a quality assessment process and read elimination (¡ Q20) using Fastp [15]. Then, a *de novo* assembly with a minimum length of 300 nucleotides was performed using MEGAHIT [16]. The contigs were compared with the non-redundant (nr) protein database of the National Center for Biotechnology Information (NCBI) with DIAMOND [17] to optimize the search for CDS regions encoding viral proteins. The files obtained were processed and analyzed with the help of MEGAN6 [17]. BLASTn and BLASTx compared the sequences of interest. To confirm the viral contigs, reads were mapped to the reference sequence of the H18N11 subtype from Peru [6] and H17N11 from Guatemala ([5] using Bowtie2 [18]. The confirmed segments were used as reference sequences to obtain segment coverage and depth with Bowtie2 and SAMTools [19]. The aligned reads were visualized and analyzed in UGENE ([20]. Finally, the genome was annotated with Prokka [21].

### Phylogenetic and evolutionary analysis

Reference genomes were obtained from GenBank. Aligning each segment’s amino acid sequences was done with MAFFT [22], and manual editing was done in UGENE. A maximum likelihood tree was then created with a bootstrap of 1000 using IQ-TREE [23] and rooted, taking into account the substitution rate measured in time with Treetime [24]. Identity patterns with reference genomes were performed with SimPlot++ [25]. Evolutionary tracking in time was executed in Bayesian Evolutionary Analysis Sampling Trees (BEAST) [26]. From the nucleotide sequences aligned in MAFFT, the substitution rate was estimated using Treetime. The evolutionary model and the options for MCMC analysis were built using the BEAUti tool, where a strict molecular clock, a GTR substitution model with *γ* distribution, and partitioning in codon positions were established. The results were visualized and analyzed in Tracer [27], where an Effective Sample Size (ESS) ¿ 200 was taken into account in all statistical analyses and the convergence of the Markov chains. Then, the information on the trees generated by BEAST was summarized using TreeAnnotator [26] and visualized in FigTree (ed.ac.uk).

### Analysis of the antigenic structure and molecular docking of the N protein

The 3D structure of the N protein of *N. albiventris* was generated by homology-based modeling with the Swiss Model [28]. The template was selected according to identity, coverage, Global Model Quality Estimation (GMQE), and Quaternary Structure Quality Estimation (QSQE). The visualization of the model was performed in ChimeraX [29]. Subsequently, the mutations concerning the N11 subtype of Peru and Bolivia were identified and mapped onto the 3D structure of the N protein of *N. albiventris*. The antigenic architecture of the N protein was then compared to the N11 crystallographic structure from Peru (4K3Y). For docking analysis, charge addition of the N protein/ *N. albiventris* and bat HLA-DR (8JRJ) was performed in ChimeraX. Then, an information-driven flexible docking approach was used for modeling biomolecular complexes using HADDOCK [30]. Van der Waals energy values, Buried Surface Area, and Z score mainly evaluated the models. Finally, the visualization and prediction of the contact points was performed in ChimeraX.

## 4 Discussion

The present work is the first study to sequence the genome of an IAV detected in *Noctilio albiventris* from Colombia. The phylogenetic analysis reveals that IAVs from *N. albiventris* formed a divergent clade in seven of the eight segments of this virus. The results are consistent with previously reported studies and provide a general overview of the process of viral adaptation to different hosts. The HA gene is the only segment related to all 16 known avian subtypes and is found within group one. These sequences share a common ancestor that gave rise to HA groups A and B, a discovery that adds a new layer of complexity to our understanding of IAV evolution.

The clade of bat IAVs, which diverged around 300-500 years ago [5], indicates that HA was initially able to mediate binding to bat receptors and then made adaptive changes to bind to MHC-II [12, 13] but maintained characteristics of primitive nodes.This finding has significant implications for the evolution and transmission of IAV. Undoubtedly, the hemagglutinin (H) of avian strains have tropism for the enteric tract due to the abundance of *α*-2,3 sialic acid that presents a “linear” structure that improves fusion with the cell, while the strains that infect humans bind to *α*-2,6 that has a “bent” structure and is mainly found in the respiratory tract [31]. However, the hemagglutinin of bat IAVs is highly divergent and binds to MCH-II [5, 6]. From this approach, the binding capacity of bat IAVs should not be analyzed based on the knowledge generated by avian IAVs [11]; instead, it should be analyzed from the phenomenon of adaptability and the ability to infect new hosts [31].

The N protein of *N. albiventris* is the most divergent among the eight segments compared to the sequences of bat and bird IAVs. This divergence suggests a new subtype called H18N12. Comparing Peru’s antigenic architecture of the N11 protein showed critical structural differences. The movement of loop 150 affected the general morphology of the protein, suggesting a structural plasticity that could influence the efficiency and specificity of the neuraminidase to perform its function during viral release [32]. The reduction in the size of the hypothetical active site implies changes in the stability of the structure and possibly in the capacity to bind to sialic acids [5, 6]. Three of the five mutations (K363R, T242K, and G361S) near the hypothetical active site increased the possibility of binding to the HLA-DR of bats. This finding generates uncertainty regarding the functions of this protein and the implications that such changes could have on the processes of viral replication or transmission. The changes generated in the protein strengthen the interactions with cellular receptors and support the theory of a down-regulation function of MHC-II by the bat IAV N protein [9]. The change of a Lys for Arg favors new interactions by increasing the number of hydrogen bonds due to the presence of a guanidino group in its side chain. Meanwhile, Met for Phe allowed the binding with hydrophobic regions of the HLA-DR of bats [33, 34]. It was shown that there is an interaction of loop 150 (Phe146), a crucial area for the enzymatic activity of this viral protein since it participates in the recognition and cleavage of receptors on cell surfaces. However, these data must be corroborated by in vitro assays.

Another aspect is the biological implications and ability of IAV to bind to MHC-II of other mammals. A recent study showed that catalytic activity could be acquired again with significant mutations in the amino acid sequence [35]. The F144C and T342A changes in the N11 protein increased viral particles in MDCK II cells compared to cells infected with the wild-type virus. Furthermore, IAV had a broader tissue tropism in the airways of mice (BALB/c) and ferrets (*Mustela putorius furo*) [35]. This is because, in the three-dimensional structure of N11, residue 144 is located at the outer edge of the hypothetical active site, while residue 342 is close to the binding site. Both sites are critical for the sialidase activity of the N protein and are present in avian IAVs [34, 36, 37]. In this sense, the zoonotic potential and interspecies jump of bat IAVs cannot be ruled out.

Studies indicate that the H17N10 and H18N11 subtypes have inefficient transmission and infection in non-bat hosts without generating critical structural changes. In the case of the IAV sequence detected in *N. albiventris*, the hemagglutinin is 100% identical to the H18 subtype. However, the neuraminidase has mutations that could improve its infection process and transmission to new species. The genetic plasticity of IAVs allows them to adapt to different species since, in the case of the bat H9N2 subtype, it has been shown that it can replicate and be transmitted between ferrets. In addition, it can efficiently infect human lung cell cultures. It can evade antiviral inhibition by MxA in B6 transgenic mice, and it also generates cross-reaction with N2-specific antibodies in human sera [38]. Bat IAVs demonstrate the ability of influenza viruses to transmit and adapt to new hosts. The new sequences must be studied to understand whether they can generate significant epidemiological outbreaks.

In conclusion, the characterization of viruses with zoonotic potential, such as the Influenza A virus (IAV) in bats, is of utmost importance as it allows us to elucidate the role of these wild animals in the evolution of the IAV virus. The virus found in our study has sequences that are new to science, with genetic and structural changes in the N protein, leading us to propose a new IAV subtype called H18N12. The mutations in the hypothetical active site of the proposed N12 subtype reveal the ability to bind to the HLA-DR of the MHC-II of bats. The detection of IAV-Bat in members of the *Noctilionidae* family expands the range of hosts where these viruses can be detected. The fishing bat *Noctilio albiventris* is associated with aquatic ecosystems, and bodies of water may be the medium of interaction between fishing bats and birds, known hosts of avian influenza. These findings underscore the need for further research to understand the potential epidemiological implications of these new sequences, and the importance of ongoing research in this field.

## 5 Author contributions statement

D.E.D contributed to the protein structure prediction, molecular docking and writing of the manuscript. D.E.D and R.R contributed to the quality control of the genome, bioinformatic and phylogenetic analyses. D.E.D, C.M.B, R.H, M.A, E.G, and V.B carried out the sample collection, RNA sequencing and data analysis. D.E.D, B.G.P, Y.L, G.A, and S.M contributed to the methodological design of the study, results analysis and discussion. All the authors reviewed and approved the manuscript.

## 6 Competing interests

The authors declare no conflict of interest.

## 7 Funding

This research was founded by a grant of the Ministry of Science, Technology and Innovation of Colombia. Code 91722-contract 601-2022. Project: “Strengthening public health research capacities: Metagenomics of infectious agents in mosquitoes and bats from four departments of the Colombian Caribbean.” R.R was supported by funding to Verena (viralemergence.org) from the U.S. National Science Foundation, including NSF BII 2021909 and NSF BII 2213854.

## Notes

### Competing Interest Statement

The authors have declared no competing interest.

### Summary of Updates

Updated document. An accidentally omitted author was added in the PDF version of the final manuscript.

